# Simera: Modelling the PCR Process to Simulate Realistic Chimera Formation

**DOI:** 10.1101/072447

**Authors:** Ben Nichols, Christopher Quince

## Abstract

Polymerase Chain Reaction (PCR) is the principal method of amplifying target DNA regions and, as such, is of great importance when performing microbial diversity studies. An unfortunate side effect of PCR is the formation of unwanted byproducts such as chimeras. The main goal of the work covered in this article is the development of an algorithm that simulates realistic chimeras for use in the evaluation of chimera detection software and for investigations into the accuracy of community structure analyses. Experimental data has helped to identify factors which may cause the formation of chimeras and has provided evidence of how influential these factors can be. This article makes use of some of this evidence in order to build a model with which to simulate the PCR process. This model helps to better explain the formation of chimeras and is therefore able to provide aid to future studies that intend to use PCR.

## 1 Introduction - Why is a New PCR Model Required?

Whilst a number of PCR models exist, there is a sparsity of models built for the purpose of artificial chimera generation. Those that do simulate chimeras, do so in such a way that the amount produced is based on the user’s desired number of chimeras. A more realistic model would rely on the composition of the input sequences and values of parameters modelling PCR conditions to drive chimera generation - the number of chimeras produced and their composition should be dependent on the input and not predetermined. Simulation software meeting these requirements would be very welcome.

An advantage of simulated data is the presence of complete information - because the input data is known then it is possible to separate the output data into chimeras and good reads with 100% accuracy. If, then, the simulation proves to be realistic enough it will be extremely useful for testing chimera detection software without the required time and expense of experimental data.

It has been claimed that the leading chimera detection tools, Perseus and UCHIME, can detect nearly all chimeras in a dataset with few false positives [14] [7] but just how confidently can these assertions be made? Both Perseus and UCHIME were tested on mock community datasets with good results, however, it would be desirable to see how the results would compare if they were tested using a dataset with a more realistic community structure, chimera frequency and chimera composition. The models formulated in this article may be used to generate *in silico* datasets designed for this purpose.

If chimera removal software does not perform as well as has been imagined then this would be cause for concern. The presence of undetected chimeras in datasets could give a false picture of community structure, likely overestimating richness and diversity levels, and would ultimately add a significant degree of uncertainty to the findings of any research that has been carried out on such data.

The findings from Fonseca et al. [10] show that chimera formation is a complicated process affected by a number of different factors such as relatedness, species diversity and nucleotide diversity. All of these factors contribute and interact to influence the formation of chimeras in ways that are difficult to understand using experimental data alone. It would, therefore, be very interesting to see whether a model designed to simulate chimera formation could help to explain how this complex system works. If a model could somehow incorporate all of these factors, then the different interactions between them could be explored and it may be possible to determine which factors have the most influence on the formation of chimeras.

There is the possibility that other, as yet unknown, factors could also contribute to the level of chimera formation. In addition to this, the amount of randomness involved is not understood. A good model of the PCR process, designed specifically with chimera formation in mind, would allow comparisons to be drawn between experimental and simulated data. This would allow improvements to be made to chimera identification and noise removal techniques.

In conclusion, there is clearly a need for a PCR model that better simulates chimera generation.

### 1.1 Existing Models of PCR

Many different studies into the simulation of PCR have been carried out in the past. Differing limitations, areas of study and goals relating to the usage of these simulations have led to varying levels of complexity and various different applications.

Some existing PCR simulators operate by selecting target regions from a set of longer genome sequences when given the primer sequences as input and return the required amplicon sequences as output. Rubin et al. [15] present such a model which is designed to investigate the production of non-targeted PCR products using a simple algorithm that matches primer sequences to suitable template DNA sequences based on a maximum mismatch threshold. The study concludes that, according to the results of the simulation, more unwanted PCR products are formed in practice than predicted by the model.

Another similar PCR simulator is *ecoPCR* [8] which takes a primer pair as command line input and makes use of the Wu-Manber algorithm [18] for pattern searching. This algorithm compares two strings and indicates whether or not the longer string contains a substring that is “approximately equal” to the shorter string. In other words, two strings are treated as identical if they are within a specified *Levenshtein distance* [12] of each of other. The Levenshtein distance is, in basic terms, a measure of the number of insertions, deletions or substitutions required to convert a given string into a target string. In the context of simulating PCR, the Wu-Manber algorithm is used to search for the optimal region of a given sequence with which to bind a primer. Output from ecoPCR includes the amplicon sequence, its length, the number of mismatches on each primer and various taxonomic information relating to the sequences.

There are also several websites which offer PCR simulation via the input of sequences and primers directly into the user’s web browser as well as changing variables relating to PCR conditions. Examples of such websites are cybertory.org [3], bioinformatics.org [2] and amnh.org [1]. The usage of these tools is generally limited to data containing fewer input sequences.

*Primer Prospector* [17], whilst not designed specifically as a PCR simulation tool, may be used in the same way as much of the software described in this section. The tool assesses the ability of a primer pair to act on a dataset of sequences and outputs statistics based on the proportion of these sequences that can be expected to amplify as well as a file containing all of the amplicons generated.

As is the case with those outlined so far in this section, the majority of available tools simulate PCR by extracting the targeted sequence fragments from the reference. They predict probable PCR products and generate statistics about potential mismatch locations and primer efficiency but they do not imitate a PCR process. An exception to this is *Grinder* [4] which produces simulated PCR amplicons with chimeras and single-base PCR errors included. Chimeras may be generated from an input parameter specifying the percentage of chimeras required and, similarly, the number of PCR errors can be controlled by inputting the required mutation rate and distribution. In Grinder, a chimera may be generated in one of two ways - the first method is randomly selecting a pair of parents and a random break point and the second is similar to the method used by CHSIM. Chimeras are then randomly added to the output data based on the required chimera proportion.

CHSIM is the name of the chimera simulation algorithm which was used to generate chimeras for the purpose of testing UCHIME [7]. The algorithm selects parent sequences which share an identical sub-sequence (*k-mer*) of given length, this k-mer is used as the crossover section between the two parents (i.e. the break point is contained somewhere within this section). Chimeras are generated at random, weighted in favour of those containing the most abundant k-mers present in the pool of potential parents. This is intended to make break points more likely between similar sequences in regions of high sequence similarity. A preset number of chimeras are generated in this way and added to the original pool of parents after each simulated round of PCR.

### 1.2 Choosing a Good Model

In order to choose a good model for any procedure, several things should be considered such as the model’s complexity as well as the parameters and input required for the model. The number of different variable parameters will impact on the model’s complexity and it may be decided that it is best to ignore certain variables in order to simplify the model. It is important to correctly identify the sources of variation that affect the process in practice and to model these realistically using appropriate methods. One example of this is the selection of appropriate probability distributions from which to draw random variables.

A good model should also be easy to implement and run quickly enough so as to be practical. The functionality of the model should be expressible in the form of an algorithm that can be implemented in code. When implementing the algorithm, compatibility with existing software and file formats (for input and output) must be taken into consideration. If large amounts of data are to be processed then it is desirable to use an algorithm that minimises the number of calculations in order to reduce the running time. Sometimes it may be better, or even necessary, to forfeit some accuracy in order to produce a faster algorithm.

Most factors that should be considered when choosing a good model will have an effect on its complexity and often a trade-off between complexity and accuracy will be necessary. A simple model is more desirable if it is as effective as more complicated models. However, if a model is oversimplified then there is a danger that its output will be unrealistic. For example, a very simple model of PCR would be to take as input the initial abundance of each DNA sequence and increase this amount based on the number of PCR rounds, such that

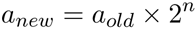

where *a*_*o1d*_ and *a*_*new*_ are, respectively, the original and resultant abundances of the sequence and n is the number of PCR rounds. To calculate the new abundance, the old abundance is multiplied by a factor of two raised to the power of *n* because each sequence splits into two new sequences during each round of PCR.

Output from this model will not be useful in practice because it does not take into account the randomness and errors inherent in PCR amplification. In particular, it ignores the facts that amplification is not 100% efficient and that the amplification step can fail before completion, creating artefacts that further complicate matters.

## 2 Methods

Chimera break point distributions taken from experimental and simulated data were compared using the *two sample Kolmogorov-Smirnov test* [13]. This test returns a p-value to indicate the probability that the two samples are similarly distributed. This means that an insignificant p-value (typically *p* > 0.05) will reveal no information about the similarity of the two sample distributions but it can be concluded that they are similar enough that there is no obvious distinction.

The Kolmogorov-Smirnov test is typically used for samples with continuous data, however it has been adapted for discrete samples in the **R** package, *dgof,* and is therefore applicable for the analysis of break point distributions. Before each Kolmogorov-Smirnov test was carried out, the larger of the two samples being tested was sub-sampled to the same size of the smaller. Because different sample selections give different p-values, the process was repeated 100 times in each case and the mean p-value was taken.

Correlation between break point frequencies occurring in experimental and simulated output was assessed using *Pearson’s correlation coefficient*, *r*_*XY*_, which is calculated using the formula,

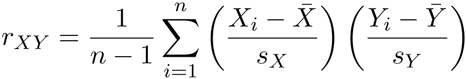

where *n* is the number of observations in each sample (both samples must contain the same number of observations), *X_i_* and *Y_i_* are the break point frequencies at position *i* in sample *X* and sample *Y* respectively, 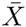 and 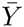 are the mean break point frequencies across all positions in sample *X* and sample *Y* respectively and *s_X_* and *s_Y_* are the sample standard deviations. As for the Kolmogorov-Smirnov test, the two samples were sub-sampled to the same size 100 times and the mean of the 100 different Pearson’s correlation coefficients was recorded.

Similarity in nucleotide composition between simulated chimeras and chimeras generated experimentally was assessed using the ‘global search’ function in *USEARCH* [6]. One dataset of chimeras (e.g. experimental chimeras) was used as a query dataset to be searched against a reference dataset (e.g. simulated chimeras). Sequences in the query dataset were paired with the most similar sequence in the reference dataset and a similarity score was recorded (number of matching nucleotides divided by alignment length).

## 3 The PCR Process

This section presents a summary of PCR as a procedure, the steps of which must be emulated to develop a realistic model of the process. The PCR process is also summarised visually in Figure 1.

**Figure 1:**
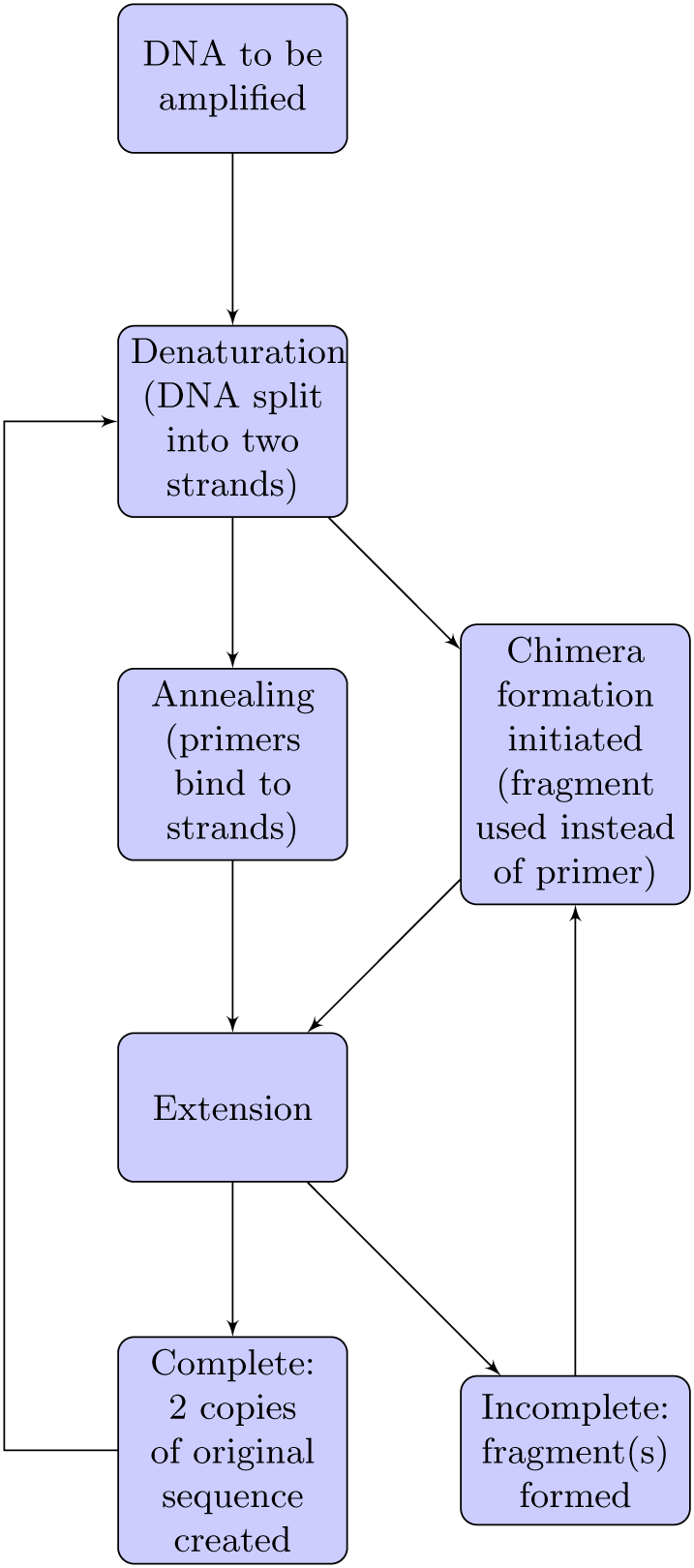
The PCR process.

In order to prepare a sample for sequencing, an amplification step is carried out using Polymerase Chain Reaction (PCR). Thermal cycling is used to repeatedly melt and cool the DNA. When a strand of DNA is copied, this copy can then also be copied; this leads to an exponential amplification effect. PCR is used to amplify a particular target region of the DNA - this is selected using primers (small pieces of DNA, complementary to the target region).

The process typically involves 20-40 cycles of the following steps (2^40^ gives approx 10^12^ copies):

1. Denaturation - this step takes place at temperatures between 94 and 98°C for around 20 to 30 seconds. Hydrogen bonds are broken to split the DNA into two strands.
2. Annealing - the temperature is reduced to 50-65°C. The primers bind to both single strands of DNA. Hydrogen bonds are only able to form when there is a close match, ensuring that the primers are annealed to the correct region.
3. Extension - the temperature is adjusted depending on the polymerase used. Nucleotides are attached to complete the DNA strands. These strands can now be copied in the same way as the original.

### Forward and Reverse Primers

After the annealing step, when the DNA molecule has been split into two strands, the primer binding onto one of these strands is called the *forward primer.* Extension can only occur in the 5’ → 3’ direction, this means that the primer binding to the second strand of the complementary pair must induce extension in the opposite direction to the first. A different primer, the *reverse primer,* must be used for this.

### Chimera Formation

Chimeras can be formed when the PCR extension step is incomplete. If PCR fails at a certain point then an incomplete sequence of DNA is produced, this fragment can act as a primer for a different sequence in another round of PCR. This has the effect of forming a sequence which is really a combination of two or more different partial sequences. The proportion of chimeras present varies from dataset to dataset. Some datasets can be comprised of 90% chimeric reads. This is obviously a large problem that is addressed using noise removal software.

## 4 Model 1

### 4.1 Model Outline

The repetitive cyclic nature of PCR suggests that an intuitive model is an iterative procedure with the same steps being repeated for every simulated round of PCR. The basic input information that will be required are the number of rounds of PCR, the primers to be used, the DNA sequences to be amplified and their initial abundances.

There are two factors that drive the way PCR progresses. The first of these is the rate of failure of PCR, when the two parts of a DNA strand do not combine with primers or fragments to begin amplification and, instead, simply recombine with each other. This failure rate will depend on the relative concentrations of sequences, primers and fragments and can be calculated each round. The second factor is the rate of failure during extension. Parameters used in this model should be chosen with these factors in mind.

One possible approach for modelling PCR, and the approach used in this article, is to use integer values for the abundance of each sequence. This means that a sequence will be treated in the model as an individual strand of DNA and allows the model to be closely analogous to the actual process. Because of this, discrete probability distributions, such as the binomial distribution, will be required to generate the random variables necessary for the model.

The steps which make up the algorithm for the model are described in detail in the remainder of this section. The complete algorithm is referred to as *Simera,* a portmanteau of the words “simulation” and “chimera”. The Simera algorithm is presented in Figure 2 and visualised in Figure 3.

**Figure 2:**
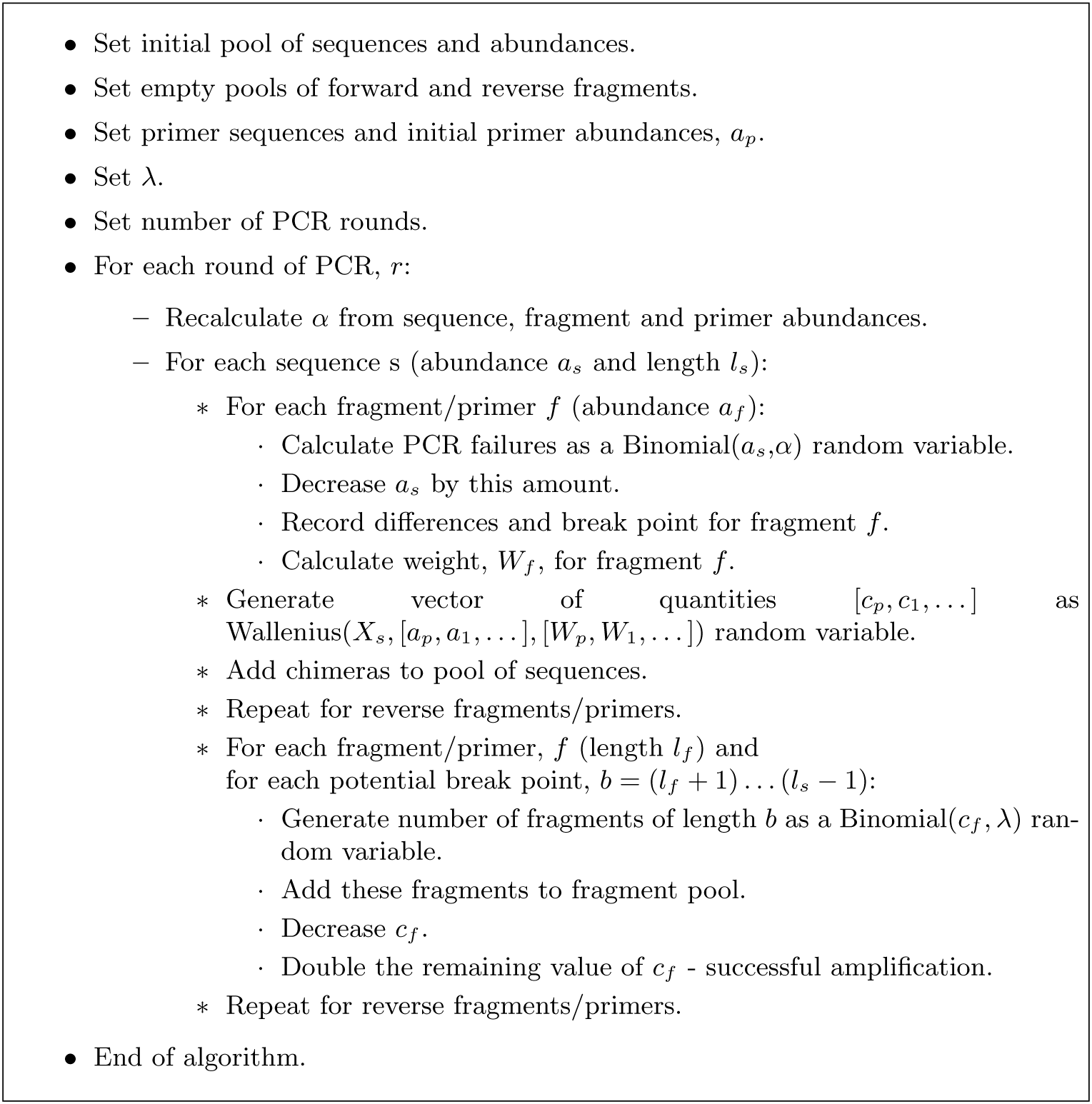
Simera algorithm for Model 1. λ is the rate of failure during PCR extension at each nucleotide point on a sequence. α is the PCR failure rate and is calculated using the formula in Section 4.1. The fragment weightings, *W_f_*, are calculated using the formula in Section 4.1.

**Figure 3:**
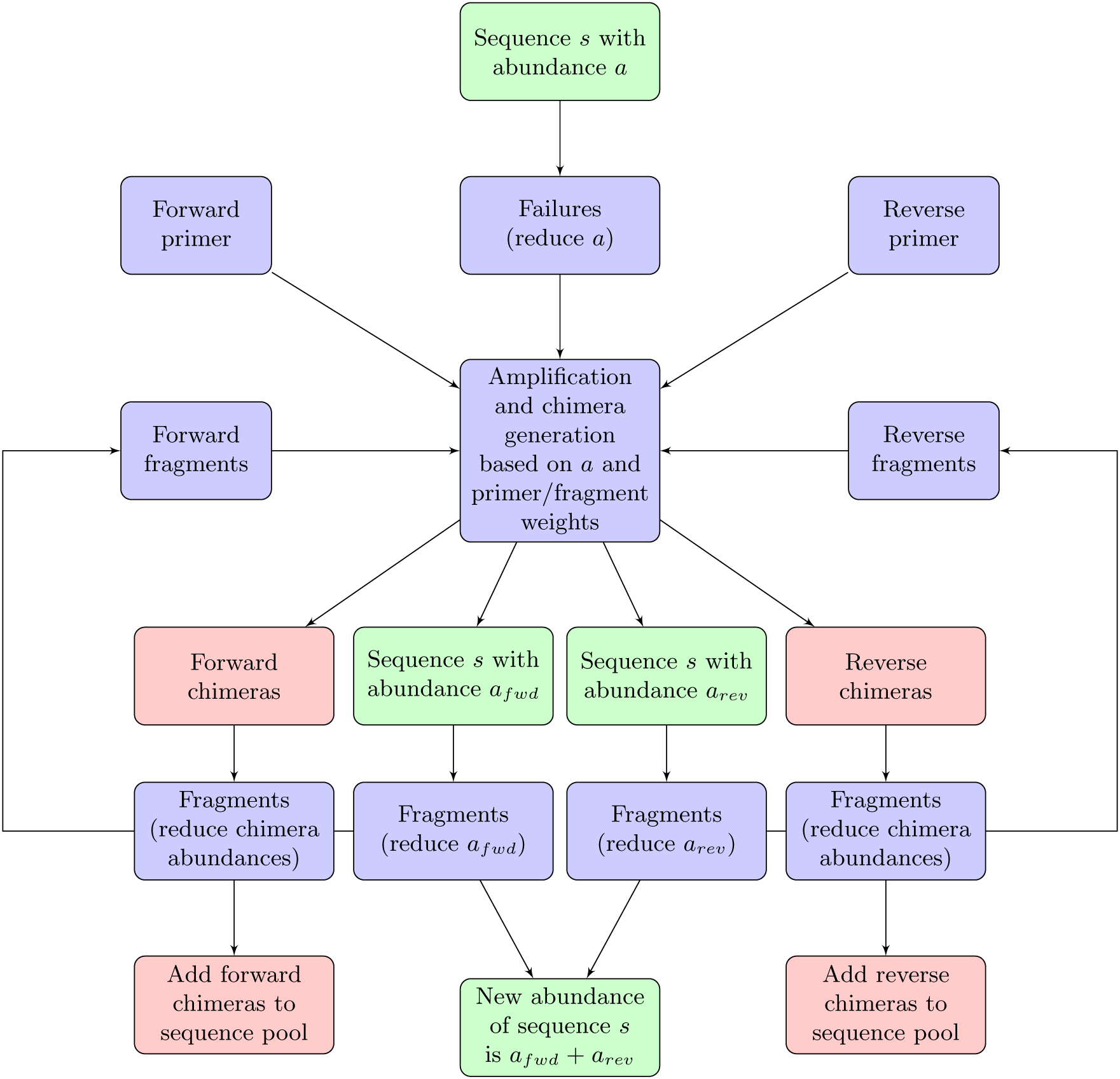
PCR simulation using Model 1. This process is performed for all sequences, s, and repeated for the desired number of PCR rounds.

#### 4.1.1 Assumptions

For this model it is assumed that failures during the PCR extension step will occur at a fixed rate. That is, extension is equally likely to fail regardless of the position on the DNA strand being amplified and regardless of the nucleotide content at this position.

Complete PCR failures - PCR failures without any extension - are assumed to be dependent on the relative primer abundance which will decrease in later rounds.

It is assumed that the ability of a fragment to act in place of a primer is directly affected by its degree of similarity to the true primer. Further to this, it is assumed that fragments generated by forward extension (5′ → 3′) may only act in place of forward primers and those generated by reverse extension (5′ ← 3′) may only act in place of reverse primers.

#### 4.1.2 Input Parameters

1. *n* - The number of rounds of PCR to be simulated.
2. λ - This parameter is the rate of failure, during the extension step, at each nucleotide on a sequence. It is used to determine if the first nucleotide is duplicated, then the second, etc. until the entire sequence has been amplified. If amplification fails at any point then a fragment is produced. λ is a probability between zero and one, and should typically be very small. λ may depend on PCR conditions so should be variable from dataset to dataset.

#### 4.1.3 Sequences

A list of initial sequences and their relative abundances shown as integer values are required as input for the Simera algorithm. The sequences will each be a string of DNA nucleotide codes [A,C,G,T] and only the region selected for amplification need be included. In practice, for implementations of the model, a fasta file is a good way to represent these data.

#### 4.1.4 Primers

Information about the forward and reverse primers must be also be supplied as input. The primers will be a string of DNA IUPAC codes [A,C,G,T] and will typically be about 20 base pairs long. Unlike the DNA sequences, primers may also contain ambiguous IUPAC codes [R,Y,S,W,K,M,B,D,H,V,N] which each represent two or more of the four specific DNA nucleotides. For example, a primer containing the code M in the first position actually represents a collection of primers where 50% contain the A nucleotide and 50% contain the C nucleotide in the first position.

These codes are included in primers because they are more versatile and can, therefore, be better at selecting sequences which have a high degree of nucleotide variation at certain points. The ambiguous IUPAC codes and the nucleotides which they represent are shown in Table 1.

As input data, the abundance of each primer is also required. This should be an integer value and should be greater than the number of primers required to perfectly amplify all sequences for the given number of rounds, *n*. Therefore, if the initial sequence abundance is *a_seq_* then the initial primer abundance should be

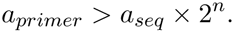

**Table 1:** Representation of specific DNA codes by ambiguous IUPAC codes. The non-zero entries show which of the four nucleotides (A,C,G,T) each IUPAC code is capable of representing.

| Code | Proportion of A | Proportion of C | Proportion of G | Proportion of T |
| --- | --- | --- | --- | --- |
| R | 1/2 | 0 | 1/2 | 0 |
| Y | 0 | 1/2 | 0 | 1/2 |
| S | 0 | 1/2 | 1/2 | 0 |
| W | 1/2 | 0 | 0 | 1/2 |
| K | 0 | 0 | 1/2 | 1/2 |
| M | 1/2 | 1/2 | 0 | 0 |
| B | 0 | 1/3 | 1/3 | 1/3 |
| D | 1/3 | 0 | 1/3 | 1/3 |
| H | 1/3 | 1/3 | 0 | 1/3 |
| V | 1/3 | 1/3 | 1/3 | 0 |
| N | 1/4 | 1/4 | 1/4 | 1/4 |

#### 4.1.5 Fragments

Two lists of sequence fragments are also required. Initially these are empty but, during the simulation, fragments will be generated and recorded. The first pool is a pool of forward fragments - those generated from forward primers - and the second is a pool of reverse fragments. The abundance of each fragment is defined the same way as in the pool of sequences. During the simulation the primer abundance and the abundance of incomplete sequence fragments will be used to calculate the probability of chimera formation where a fragment is selected in place of the primer.

#### 4.1.6 PCR Failure

The first step in the Simera algorithm is to calculate how many copies of each sequence fail to amplify. These sequences are determined at the beginning of each round, and their numbers are reduced accordingly so that the inactive sequences are not referenced during the amplification step.

This will be dependent on the ratio of total sequence abundance to total combined sequence, primer and fragment abundance - i.e. the fraction of all elements present in PCR that are comprised of full sequences. In the first round of PCR there will be relatively many primers (but no fragments) and few sequences so this ratio will be small. As the rounds progress, more sequences will be generated and primers will be used up so the ratio will increase in size. It is logical to conclude that if primers and fragments are in plentiful supply then there will be fewer instances when sequences fail to bond with them to instigate amplification. This reasoning has been confirmed from results that show PCR efficiency is at its highest when amplicon quantity is at its lowest and vice versa [16].

To determine how many sequences fail to amplify completely in each round, the PCR failure rate is calculated as the parameter *a* and used to generate a binomial random variable for each sequence:

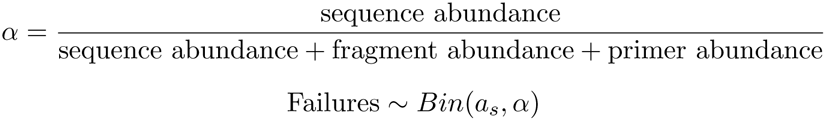

where *a_s_* is the abundance of sequence *s*. The effective abundance of sequence *s*, *X_s_* is the remaining number of molecules of sequence *s* that are available for PCR extension and chimera formation.

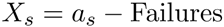

#### 4.1.7 Dealing With Reverse Primers

If the model is to follow the PCR process analogous then, when the simulation of a sequence splitting into two strands takes place, two differing sequences should be recorded. The first will be the original sequence of nucleotides and bind with the forward primer before commencing extension. The second sequence will be the *reverse complement* - meaning that the order of the sequence is reversed and that each nucleotide is swapped for its corresponding complementary nucleotide (A ⇔ T and C ⇔ G) - of the first sequence and will bind with the reverse primer.

In order to increase efficiency (and conserve memory in the implementation of the algorithm) a good shortcut is to use the reverse compeiment of the reverse primer instead of the genuine reverse primer. This means that both complementary strands for every sequence do not need to be recorded and instead only one strand is required. Binding can be simulated by attaching the forward primer and the new (reverse complement) reverse primer to opposite ends of two copies of this strand as shown in Figure 4.

**Figure 4:**
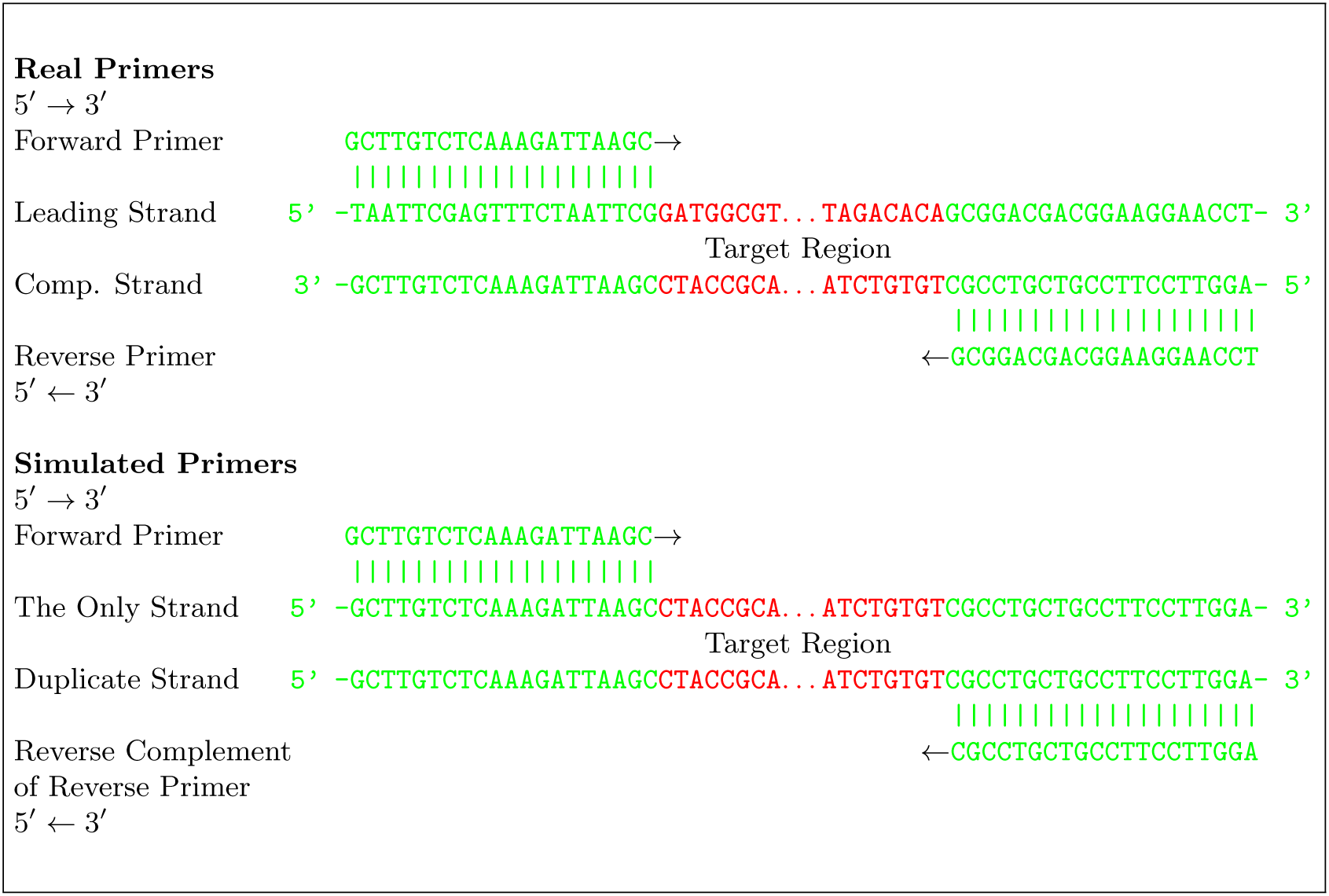
Simulated forward and reverse PCR primers. Notation referring to the direction of each primer is relative to the leading strand.

#### 4.1.8 Choosing the Best Fragments for Chimera Formation

As declared in the assumptions in Section 4.1, fragments will be less effective at binding with sequences than primers so, to make the model realistic, fragments must be penalised by giving more weight to the probability of a sequence binding with a primer. Some fragments will also be more adept than others at acting as primers so this must also be taken into account. This is done by comparing the last twenty nucleotides on the candidate fragment with all possible positions on the sequence. The functional part of a typical PCR primer is around twenty nucleotides long, therefore using twenty nucleotides from a fragment is a logical choice when the fragment will be acting as a primer.

The number of differences between the fragment and the sequence at each point is recorded, giving the minimum number of differences and the position at which this minimum value occurs for each candidate sequence.

In the example in Figure 5 it can be seen that position C gives the fewest differences between the fragment and the sequence. In this case there are zero differences compared to two and one in positions A and B respectively. So far, only fragments acting as forward primers have been considered. Fragments acting as reverse primers are analysed separately in the same way, except that the *first* twenty nucleotides of the fragment are compared with the sequences instead of the last twenty.

**Figure 5:**
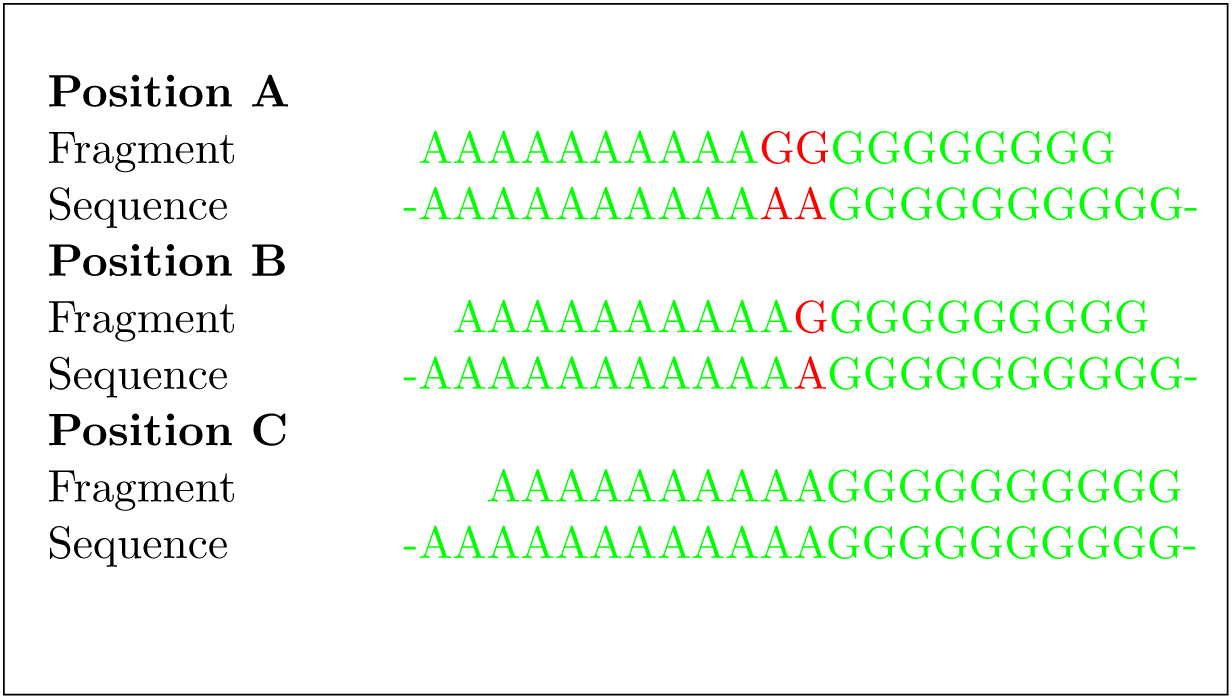
Determining the optimal position for a fragment to act as a primer. Position C is chosen because there are fewer differences.

Once the optimal position and the number of differences has been found for each fragment then each fragment f can be weighted based on its suitability using the set of parameters

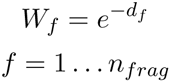

where *d_f_* is the number of differences for fragment *f* and *n_frag_* is the total number of fragments. This assigns higher weights to fragments with fewer differences, as required and all weights are forced to be between zero and one. The equation for *W_f_* takes into account the fact that a greater quantity of energy will be required to bind fragments with a large number of differences, making it much less likely that these fragments will successfully bind.

A weight, *W_p_*, for selecting a primer is calculated in the same way. In the case where the primer contains ambiguous IUPAC codes, a non integer number of differences may be awarded if parts of the primer result in a partial match to the sequence. For example, if the primer contains the code M then this will result in a difference of 0.5 if it is compared with either of the codes A or C (see Table 1). The primer is designed to be able to align well with part of the sequence so it will, typically, have very few or zero differences. It is easy to see that if there are zero differences between the primer and part of the sequence then a value of *W_p_* = 1 will be calculated.

These weights, together with the set of abundances of each primer and fragment, can be used to determine which primer or fragment each sequence will bind with. Wallenius’ multivariate non-central hypergeometric distribution can be used for this purpose because it models the selection of items without replacement based on their abundance and allowing unequal probabilities of selecting items of differing type, such as the primers and fragments of varying quality in this model. Selection without replacement is appropriate because when a primer or fragment binds with a sequence then it will no longer be available for use in the current round.

For each sequence, random variables are drawn from the Wallenius distribution to identify the quantity of each primer or fragment to be selected for amplification.

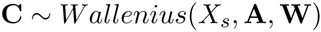

where

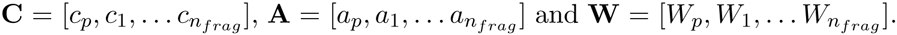

The parameters *a_p_* and *a_1_* … *a_n_frag__* are the abundances of the primer and fragments respectively.

#### 4.1.9 Amplification and Fragmentation

Each sequence can either bind with a fragment to form a chimera or bind with the correct primer to commence amplification. Amplification can either continue until the entire sequence has been amplified as intended or it can fail part of the way through to form a sequence fragment. When a sequence is ready for amplification it will be split into two strands, one will use the forward primer (or a forward fragment) and the other will use the reverse primer (or a reverse fragment). This means that amplification can be split into two separate processes. For each sequence to be amplified the abundance is set to zero then increased by one if the forward strand successfully amplifies and increased by one again if the reverse strand amplifies.

Consider the process to amplify forward strands - the reverse process is symmetrical and will not be described in detail. The sequences can be examined in turn. The parameter λ is used to determine whether the first nucleotide in the sequence is amplified. If it is then the second is amplified with the same probability and so on until the entire sequence is amplified. If at some point a nucleotide fails to amplify then amplification stops entirely for the sequence and the incomplete sequence is added to the pool of (forward) fragments. To model this, the primer and fragments are examined separately and binomial random variables are used for each possible fragment of sequence *s*. In the case of the primer,

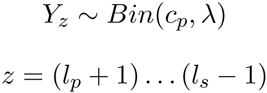

where *l_s_* and *l_p_* are the length of the sequence and the primer respectively. The new fragment is created by joining together the *l_p_* nucleotides of the primer with nucleotides in positions (*l_p_* + 1) to *z* in sequence *s*. *Y_z_* copies of the fragment are added to the pool of fragments and *c_p_* is reduced by *Y_z_*. The process is then repeated for each (old) fragment, *f*, in place of the primer, substituting *c_f_* and *l_f_* for *c_p_* and *l_p_*, respectively.

For each sequence there are (*l_s_* − *l_p_*) possible fragments. This is the case because amplification can fail at position (*l_p_* + 1) through to position *l_s_*, giving (*l_s_* − *l_p_*) possible failure points. The integer *z* is the same as the length of the fragment created.

The PCR round is now complete and a new round can commence.

### 4.2 Implementation

The Simera algorithm was implemented using C++ code. This implementation makes use of the *randomc* and *stocc* libraries [9] which provide the random number generator and probability distributions necessary to implement the algorithm. The latter of these libraries required slight modification to enable compatibility.

The program requires as input the sequences to be amplified and their initial abundances, the primer pair, the number of rounds of PCR to be simulated, the number of reads to be sampled post-simulation and the value of the parameter λ.

#### 4.2.1 Pre-processing and Formatting

The input files and parameters must be in the correct format for the Simera program to function correctly. The number of rounds of PCR to be simulated, the number of reads to be sampled post-simulation and the value of the parameter λ can be supplied as command line input and the primer pair can be supplied as a fasta file. The sequences to be amplified must also be in fasta format with each sequence having a unique name containing the sequence’s abundance as the final part of this name. The sequences themselves must be truncated so that only the target regions to be amplified, flanked on either side by the two primer-compatible regions, are present.

### 4.3 Calibration

To determine the value of the parameter λ, simulated data were compared with the experimental data described by Fonseca et al. [10]. To mimic this experiment, the good sequences (as detected by Perseus) from the experiments containing 12, 24 and 48 closely and distantly related nematode species were used as input for 35 simulated rounds of PCR - the same number of rounds as the original experiment. The same number of reads produced for each experiment were sampled from the output of each corresponding simulation and the number of chimeras produced in each case were recorded. Different values for λ were tried, each experiment was repeated five times and the value that gave the closest match between the experimental data and the simulated data was found to be λ ≈ 5 × 10^−6^ as can be seen in Figure 6.

**Figure 6:**
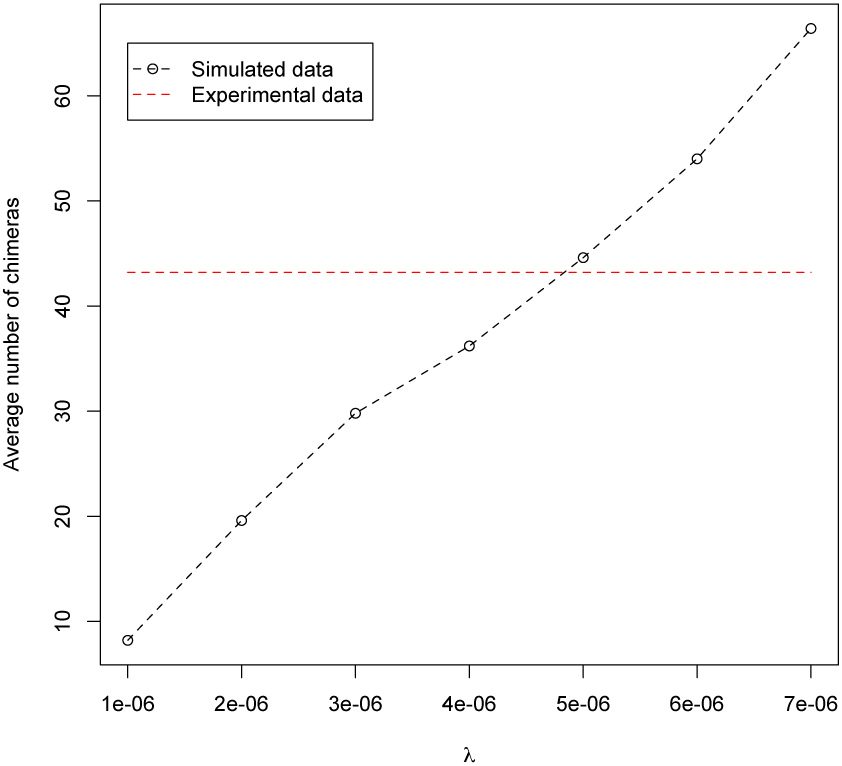
Number of chimeras simulated using the Simera algorithm for different values of λ. The parameter λ is the failure rate, during the extension step, at each nucleotide position on the query sequence. 35 rounds of PCR were simulated using the good sequences (as detected by Perseus) from pooled experiments on 12, 24 and 48 closely and distantly related nematodes as input.

This value for λ can be considered accurate for simulations of PCR under the same conditions as those used to generate the experimental data. To simulate PCR with different conditions, different values for λ may be more appropriate.

### 4.4 Results

In order to assess the performance of the model, simulated data were again compared with the experimental data described by Fonseca et al. [10]. The good sequences (as detected by Perseus) from each of the closely and distantly related pooled nematode experiments were used as input for 35 simulated rounds of PCR. True break points and parents are available as output from the simulation software, however to compare the simulated data with realistic data it was necessary to find the break points and parents in the same way as the original experiment.

As with the analysis described by Fonseca et al. [10], the output from Perseus returned most likely break point for each chimera based on its two identified parent sequences. These break points were standardised for the whole dataset by forming a four-way alignment of each chimera, its two parents and the *C. elegans* reference sequence using ClustalX [11]. The position of each break point on the reference sequence was recorded to give a standardised break point. The frequency of each standardised break point could then be recorded to assess which regions of a sequence were most susceptible to chimera formation.

Break point frequencies from the experimental and simulated data are shown in conjunction with the equivalent results for the second algorithm in Section 5.4 (Figure 12) and their distributions appear to be similar. A Kolmogorov-Smirnov test, adapted for use with discrete distributions in the *dgof* **R** package [5], was performed and yielded a p-value of 0.607, indicating that there was no evidence that the two sets of data were drawn from distinct distributions. In addition to this, the two sets of break point frequencies have a Pearson’s correlation coefficient of 0.735. It can be inferred from these results that the simulated data are distributed similarly to the experimental data.

## 5 Model 2

### 5.1 How Can Model 1 be Improved?

It has been shown in Section 4 that the first PCR model is a faithful model of the PCR process which can accurately simulate the generation of realistic chimeras. The main negative issue with the model is that the implementations of it run too slowly to be useful for studies involving medium-sized to large datasets.

Ways of generalising and adapting the model must, therefore, be sought in order to increase the speed of simulations without significantly reducing the accuracy and reliability of the output. This is achieved in this section by taking a more abstract approach which involves creating a pool of the most likely chimeras prior to the main body of the algorithm being executed. All simulated chimeras may now only come from this pool and this, in turn, means that individual fragments no longer need to be recorded. Instead, only the overall number of fragments is required.

### 5.2 Model Outline

The updated algorithm for Model 2 is named *Simera 2.* The two parts of the Simera 2 algorithm are described in detail in the remainder of this section. The complete algorithm is presented in Figure 7 and visualised in Figures 8 and 9.

**Figure 7:**
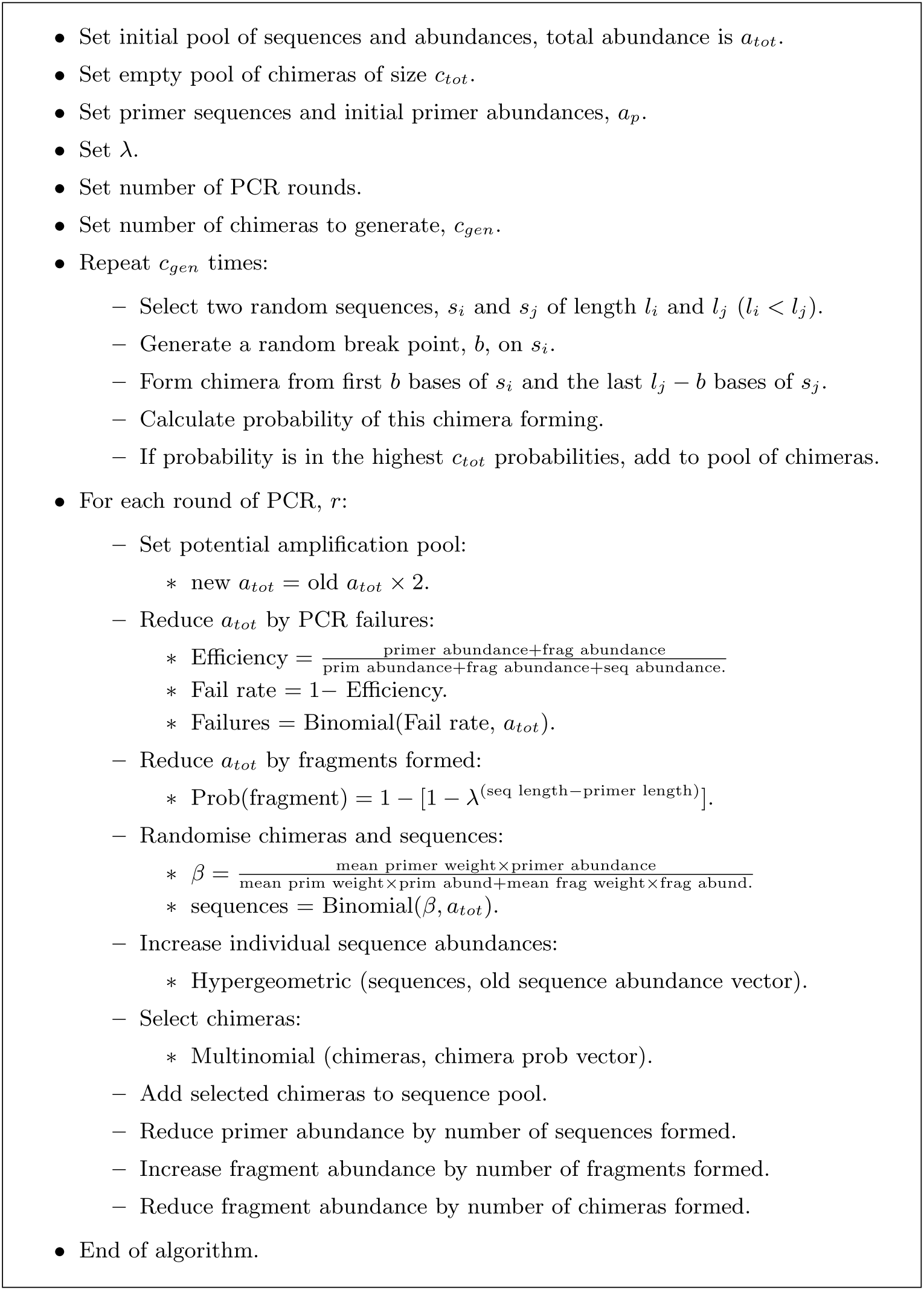
Simera 2 algorithm for Model 2. A is the rate of failure during PCR extension at each nucleotide point on a sequence.

**Figure 8:**
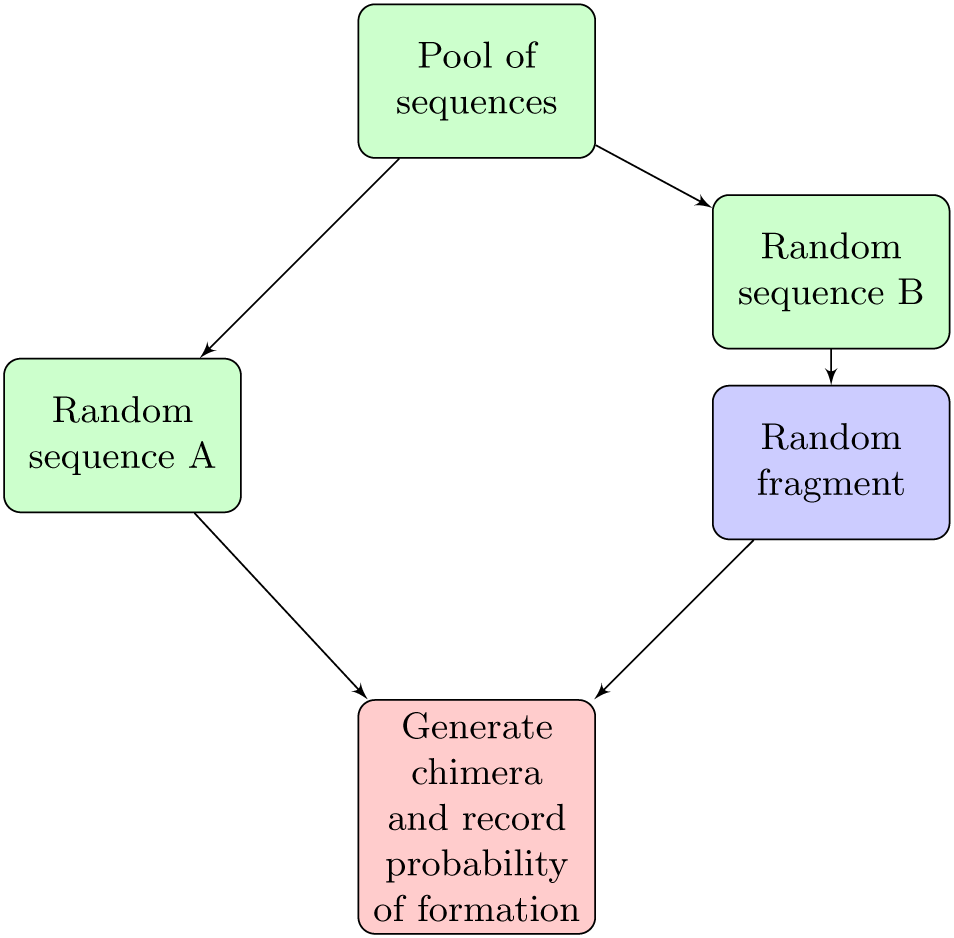
PCR simulation using Model 2: Chimera formation step. This step is to be repeated until the desired number of chimeras is reached.

**Figure 9:**
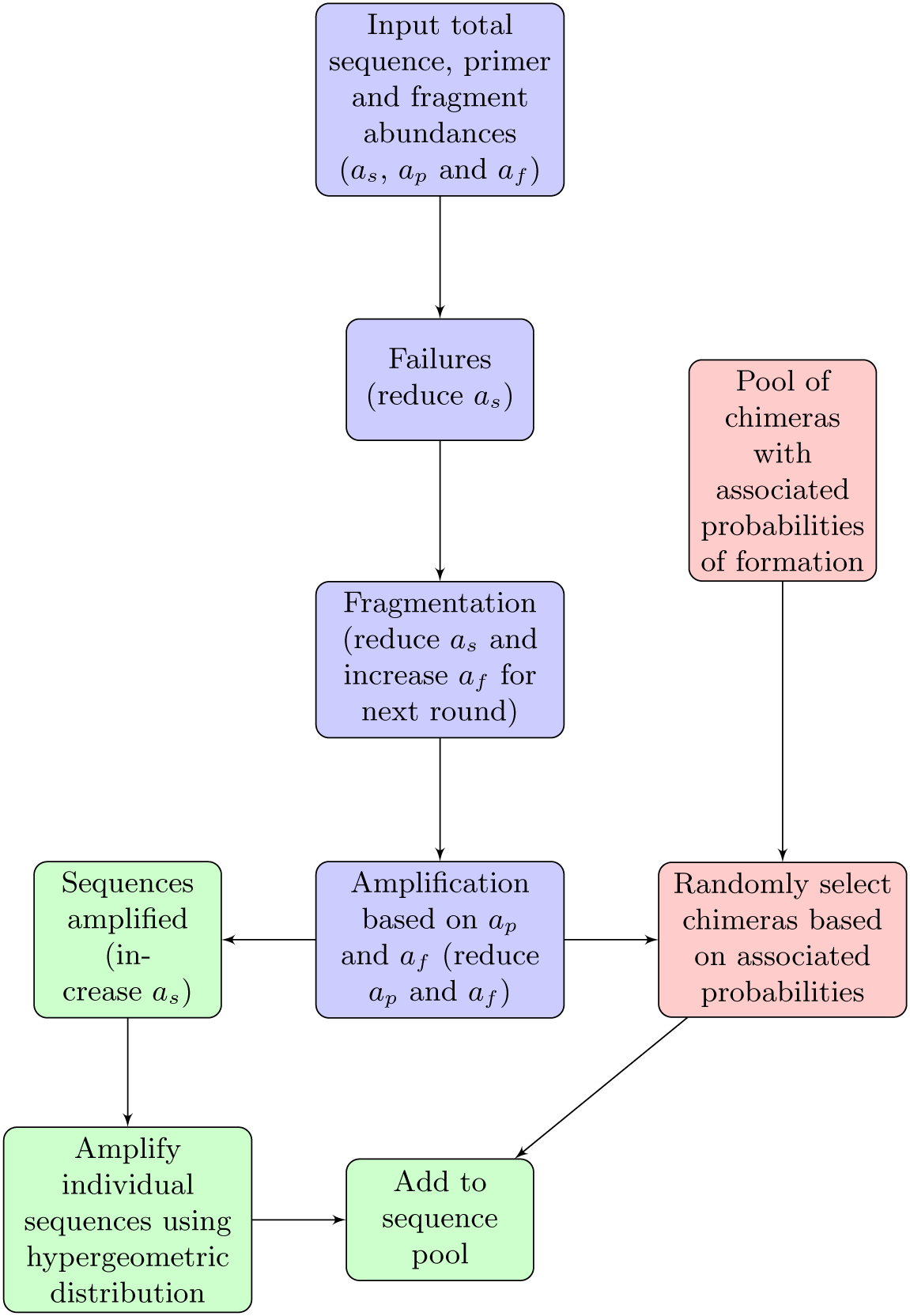
PCR simulation using Model 2: PCR step. Repeat this step for the desired number of PCR rounds.

#### 5.2.1 Assumptions

In this model chimeras are still formed in the same way, the difference is that rarer chimeras will now be ignored. Therefore, in addition to the assumptions made for Model 1, it is assumed that rare chimeras will be generated in low enough abundances during PCR that they will not be selected when reads are sampled during sequencing.

#### 5.2.2 Input and Parameters

Most of the input for the second algorithm is the same as the first:

1. A list of initial sequences and their initial abundances.
2. Forward and reverse primers and their initial abundances.
3. λ - The rate of failure at each nucleotide on a sequence.
4. A list of potential chimeras - a specified number of chimeras are to be recorded for use later on in the algorithm. The pool is empty initially.

#### 5.2.3 Chimera Formation Step

The Simera 2 algorithm comprises of two steps. The first of these involves constructing all possible chimeras and calculating the probability of each chimera forming. At a later stage in the algorithm, the probabilities associated with the individual chimeras will be used to select a chimera at random when one is created. This removes the need to generate a new chimera every time one is formed and, it is hoped, will not impact on the realism of the first model.

To generate all possible chimeras, all possible fragments are aligned with all sequences and the best chimera (fewest mismatches) is found, as described in Section 4.1.8. The probability of a fragment of length *l* forming is

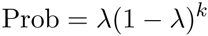

where *k* = *l* − *p* and *p* is the length of the primer used to form the fragment. The integer, *k*, is the same value as the number of successful nucleotide extensions prior to failure.

This probability is then combined with the relative abundance of each sequence and the number of mismatches between the fragment and the sequence to calculate the probability of each chimera forming,

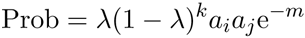

where *a_i_* and *a_j_* are the relative abundances of the two sequences and *m* is the number of mismatches.

A specified number of the most probable chimeras are recorded for later use. For larger datasets, generating all possible chimeras will be too computationally intensive and, instead, a predefined large number of chimeras may be generated randomly and the most probable chimeras are selected from these.

#### 5.2.4 PCR Step

The following PCR step is intended to approximate the first Simera algorithm outlined in Section 4, it is to be repeated for the specified number of rounds.

PCR failures are calculated using exactly the same method as in the original model, except they can be calculated for all sequences together rather than each sequence separately:

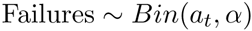

where *a_t_* is the total sequence abundance. The probability of a sequence fragmenting is

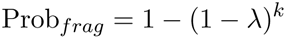

which is just one minus the probability of the sequence amplifying successfully. This probability can then be used to calculate the number of fragments created for each sequence in the current round:

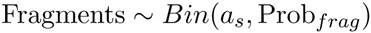

where *a_s_* is the abundance of sequence *s*. This works the same way as fragmentation in the first Simera algorithm but here only the number of fragments is recorded instead of each fragment being recorded individually. This method avoids the need to use Wallenius’ distribution to select individual fragments based on their weightings.

The number of sequences available for amplification, *a_t_*, is reduced by the number of failures and fragmented sequences.

The number of successfully amplified sequences is determined using the parameter β, where

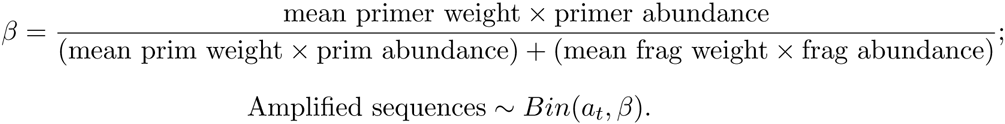

The number of individual sequences amplified is then found using a hypergeometric random variable, as follows:

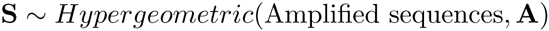

where **A** is the vector of individual sequence abundances. This amplifies the sequences all at once, compared with the first Simera algorithm which amplifies each sequence separately using successive binomial random variables. Accuracy is lost because mean fragment weights are used rather than the exact values.

Any remaining sequences are used to generate chimeras and this stage is where the two algorithms differ the most. In the Simera 2 algorithm, the chimeras are chosen from the prepared pool using a multinomial distribution rather than being created from a fragment pool when required. This is much quicker.

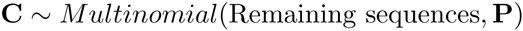

where **P** is the vector of the probabilities of formation for each chimera. Generated chimeras are then added to the pool of sequences for the next PCR round.

### 5.3 Implementation

The Simera 2 algorithm was implemented using **C++** and the program has the same input requirements and dependencies as the original Simera program.

### 5.4 Results

The implementation of the Simera 2 algorithm has been shown to be able to handle large, realistic datasets. To verify that this algorithm is a good approximation of Simera, both algorithm implementations were used to simulate chimeras for the same datasets - the closely and distantly related nematode pools which were also used in Section 4.4 - with the same input parameters (35 rounds of PCR and λ = 5 × 10^−6^). It was not possible to compare the models using larger datasets because of the inhibitive speed of the original Simera implementation. However, it was theorised that if the models work comparably on smaller datasets then the same should also be the case for larger datasets.

When the simulated output from the Simera 2 algorithm was subsampled, using the same sample size as that used for the original Simera data, the average number of chimeras produced was 42.2, compared with 44.6 for the original Simera data and 43.2 for the real experimental data.

To compare the type of chimeras which were formed, the break points were again observed. On this occasion there was no need to use the method involving Perseus and the *C. elegans* reference sequence because all break points were available as output from the simulation software. The break points of all chimeras generated in each of the simulated experiments were compared, and the distributions of these are shown in Figure 10.

**Figure 10:**
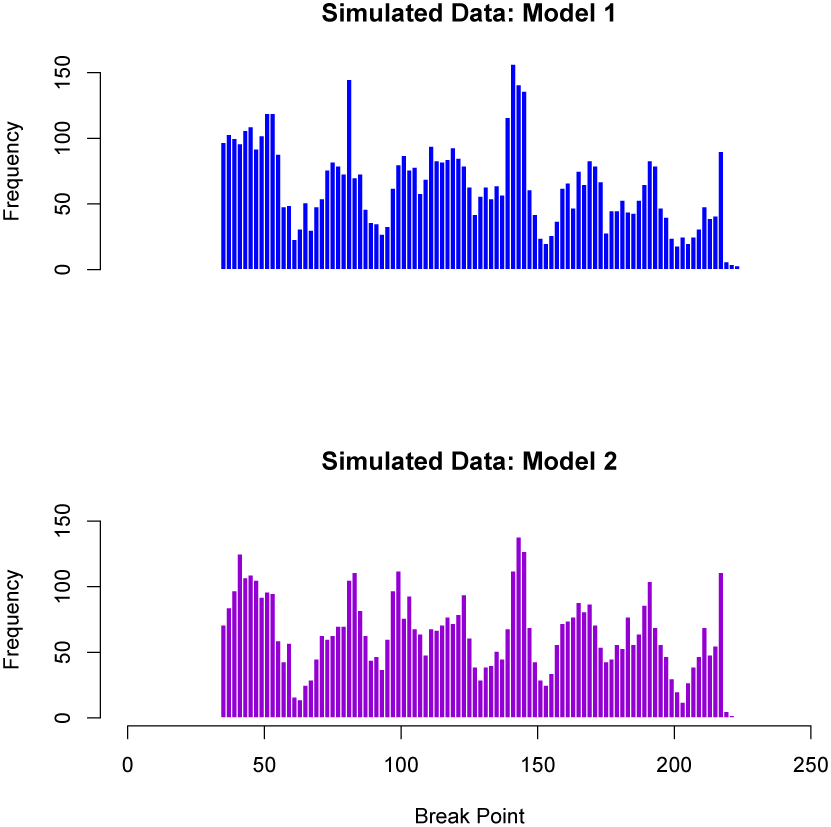
Break point frequencies for simulated data comparing results from the Simera and Simera 2 algorithms. For each algorithm, 35 rounds of PCR were simulated using the good sequences (as identified by Perseus) from pooled experiments on 12, 24 and 48 closely and distantly related nematodes as input. Break points are returned as output from the Simera and Simera 2 implementations.

The two distributions appear very alike, suggesting that both models produce the same chimeras and that Model 2 is an excellent approximation of Model 1. To verify these assertions, a Kolmogorov-Smirnov test, again using the *dgof* package in **R**, was performed on the two sets of frequency data. A p-value of 0.970 was returned, indicating that there was no evidence that the two datasets were drawn from distinct distributions. Additionally, the two datasets of break point frequencies are closely correlated, as can be seen in Figure 11, with a Pearson’s correlation coefficient of 0.914. These results provide strong evidence that the two algorithms generate very similar output when provided with identical input.

**Figure 11:**
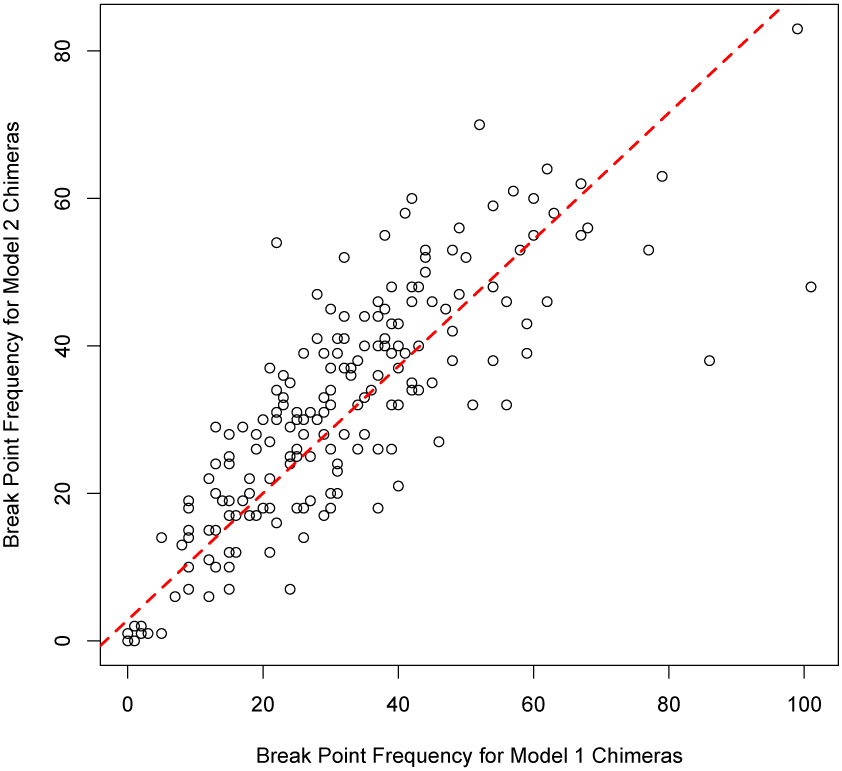
Break point frequencies for simulated data generated using the Simera algorithm plotted against the same data generated using the Simera 2 algorithm. 35 rounds of PCR were simulated using the good sequences (as identified by Perseus) from pooled experiments on 12, 24 and 48 closely and distantly related nematodes as input. Break points are returned as output from the Simera and Simera 2 implementations.

In addition to the true break points provided by the software, the break points found by aligning each chimera, its parents (as detected by Perseus) and the *C. elegans* reference sequence were also recorded and again compared with those from the experimental data. The break point frequencies are shown in Figure 12 and they appear to be distributed similarly to the experimental break points as well as the break points generated using the Simera 1 algorithm. A two sample Kolmogorov-Smirnov test between the Simera 2 break points and the experimental break points returned a p-value of 0.592, meaning that there was no evidence to suggest that the two samples were differently distributed, and the two samples were positively correlated with a Pearson’s correlation coefficient of 0.682.

**Figure 12:**
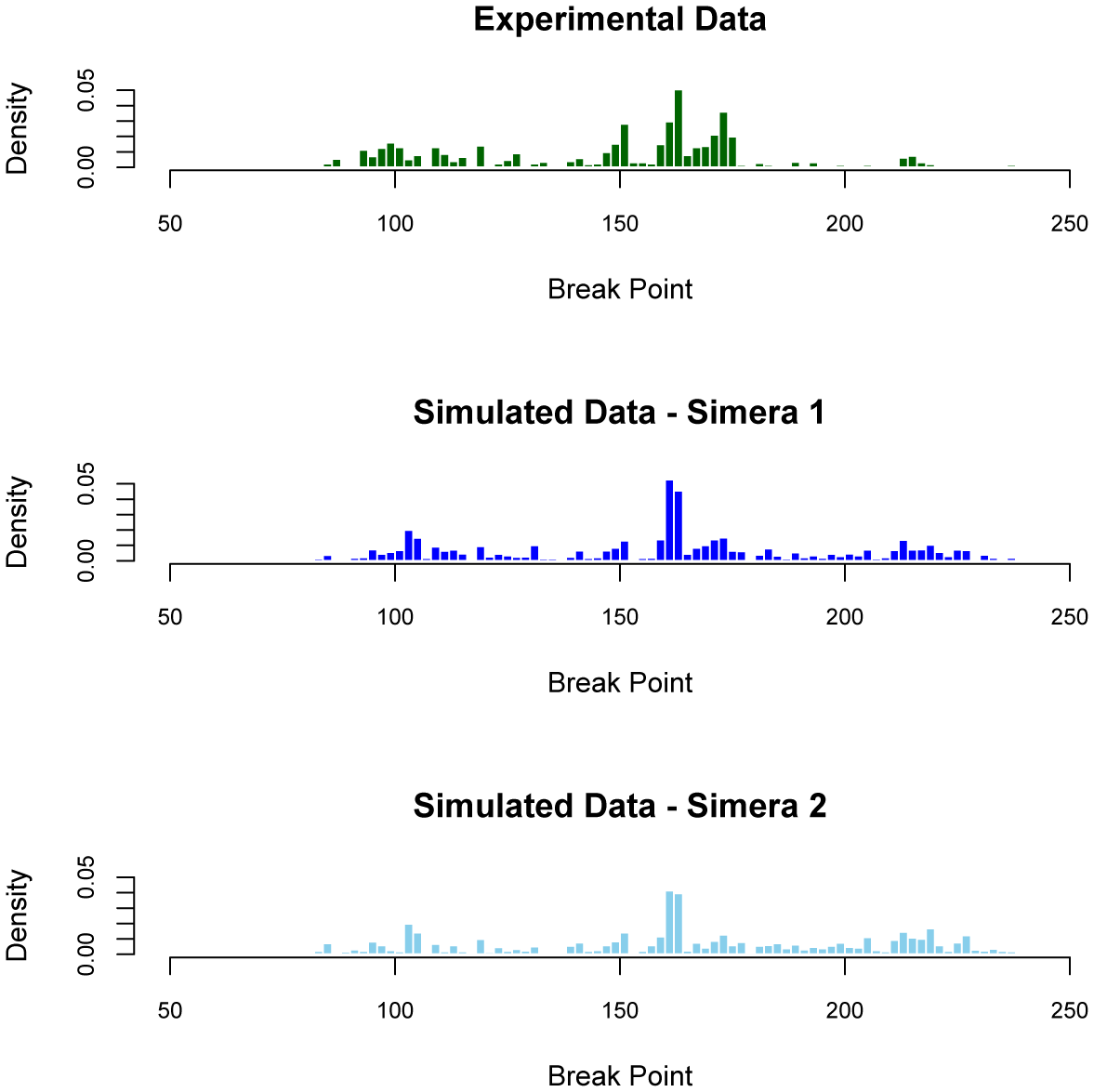
Break points of chimeras generated from the Simera and Simera 2 algorithms compared with those from pooled experiments on 12, 24 and 48 closely and distantly related nematodes. For the simulated chimeras, 35 rounds of PCR were simulated using the good sequences (as identified by Perseus) from the pooled experiments as input. In all three cases a four way alignment was formed, using ClustalX, between each chimera, its two parents (as identified by Perseus) and the *C. elegans* reference sequence. The number of break points (as identified by Perseus) at each point on the alignment were recorded.

The quality of chimeras generated with the Simera algorithms was compared with that of those generated using the existing PCR simulator, Grinder 0.5.3. Two different methods of chimera generation were used. The first was Grinder’s default method which simply creates chimeras based on a random break point, the second method applies the same technique as used by CHSIM which requires both parents to share an identical k-mer of length 10. In order to specify the required k-mer length, Grinder must be supplied with the input parameter, *‘ck’,* so the first method has *ck* = 0 and the second has *ck* = 10.

Break points for the chimeras generated by Grinder were found using the same method as was used to find the break points of those generated using the Simera algorithms, and the distributions are shown in Figure 13. As expected, the distribution when ck = 0 is fairly flat and bears little similarity to the distribution of the experimental break points. The distribution when ck = 10 seems more realistic with some peaks and troughs appearing in the same regions. However, there is an excessively large number of break points over 200 and the remainder of the peaks are not as high as their experimental counterparts. Overall, to the naked eye, the distribution does not seem as realistic as those generated from the Simera algorithms.

**Figure 13:**
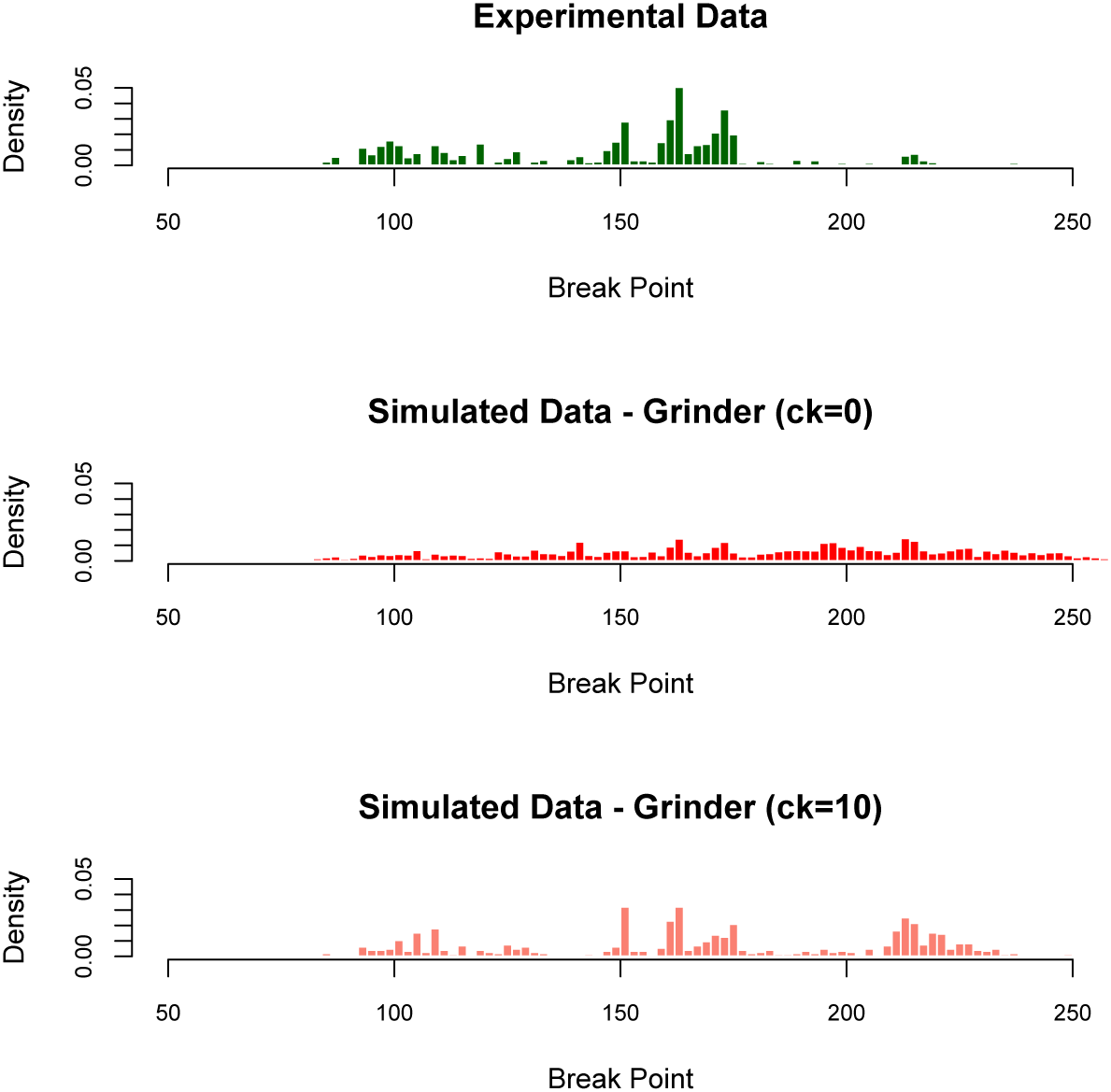
Break points of chimeras generated from Grinder, using both *ck* = 0 and *ck* = 10, compared with those from pooled experiments on 12, 24 and 48 closely and distantly related nematodes. For the simulated chimeras, 35 rounds of PCR were simulated using the good sequences (as identified by Perseus) from the pooled experiments as input. In all three cases a four way alignment was formed, using ClustalX, between each chimera, its two parents (as identified by Perseus) and the *C. elegans* reference sequence. The number of break points (as identified by Perseus) at each point on the alignment were recorded.

All simulated sets of break points were compared with the set of experimental break points and the Kolmogorov-Smirnov p-values, along with the Pearson's correlation coefficients, are displayed in Table 2. The p-value of 0.471 and correlation coefficient of 0.520 give no indication that chimera break points generated using Grinder with ck = 10 are distributed differently from the experimental data but these numbers are lower than those found using the Simera algorithms which means that there can be greater confidence that the Simera-generate chimera break points share the experimental distribution. The very low p-value shown supplies very strong evidence that the break points generated using Grinder with *ck* = 0 are not distributed in the same way as the experimental break points.

**Table 2:**
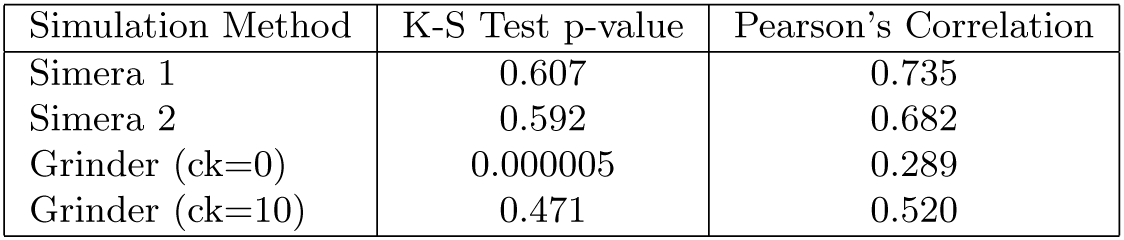
Kolmogorov-Smirnov test p-values and Pearson’s correlation coefficients returned when various sets of simulated break points were compared with experimental break points.

| Simulation Method | K-S Test p-value | Pearson's Correlation |
| --- | --- | --- |
| Simera 1 | 0.607 | 0.735 |
| Simera 2 | 0.592 | 0.682 |
| Grinder (ck=0) | 0.000005 | 0.289 |
| Grinder (ck=10) | 0.471 | 0.520 |

To investigate sequence similarity between simulated and real data, the chimeras generated from the closely and distantly related experiments were compared with the chimeras generated from the corresponding simulations using both algorithms. Using the good sequences which were detected by using Perseus on the experimental output, each simulation was repeated five times and the generated chimeras were pooled to create reference datasets for each simulated experiment. The reference datasets were subsampled so that both Simera and Simera 2 reference datasets were the same size for each experiment. USEARCH was used to find the closest matches between chimeras in the query dataset and those in the reference datasets and the number of exact matches (100% similarity) and matches with greater than 99% similarity were recorded. The two simulated datasets were compared against each other in the same way to see how many chimeras were present in both.

Figure 14 shows the results from this analysis. The output from Simera contained slightly more identical matches to the experimental chimeras than the Simera 2 output (31.5% versus 28.8%) and also slightly more near (> 99%) matches (59.7% versus 53.2%). The two simulations were shown to be producing similar chimeras with approximately 80% of the chimeras produced using Simera 2 closely matching (> 99%) those produced by Simera.

**Figure 14:**
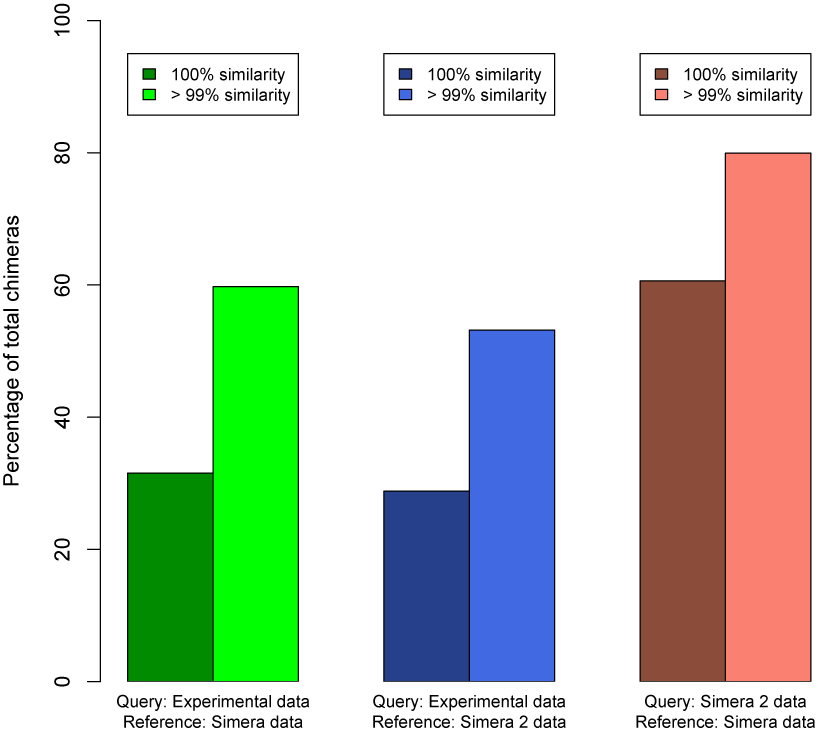
Sequence similarities (using USEARCH) when comparing experimental chimeras against datasets of simulated chimeras and when comparing datasets generated using the two different Simera algorithms.

The number of exact matches between experimental chimeras and those simulated for the reference sets was far greater than would be expected for randomly generated chimeras. Examining, for example, the experiment on 24 closely related nematodes, the number of different potential chimeras can be calculated. Each of the 24 input sequences is 220 base pairs in length, meaning that there are 200 potential break points on each sequence. This results in 200 different potential fragments for each sequence which, multiplying by 24, makes 4800 total potential fragments. Each of these fragments can form one chimera when paired with any of the 23 other sequences, so there are 23 × 4800 = 110400 possible chimeras resulting from this dataset. As there were only 7309 chimeras generated for the reference datasets used for this experiment then, under a random uniform model of chimera generation, the expected proportion of the experimental chimeras appearing in the reference datasets is 7309/110400 = 0.066 or 6.6%. This result contrasts with the actual percentage of matches for this experiment which were 38.1% for the reference dataset of chimeras generated using the Simera algorithm and 42.9% for the reference dataset of chimeras generated from the Simera 2 algorithm.

Table 3 shows the expected exact matches for each of the six experiments. In every instance these are significantly lower than the actual exact matches which is strong evidence that the simulated chimeras are more realistic than uniform randomly generated chimeras. Expected matches are lower for experiments with a higher number of input sequences because there are more potential chimeras for these. This analysis does not consider chimeras generated from other chimeras but the inclusion of these would further increase the number of different potential chimeras and, therefore, further decrease the amount of expected matches.

**Table 3:** Expected 100% matches for experimental chimeras versus actual 100% matches when compared with reference datasets of chimeras generated using the Simera and Simera 2 algorithms. USEARCH was used to determine percentage similarity between sequences.

| Species Relatedness | Number of Species | Number of Chimeras in Reference Databases | Expected 100% Matches | 100% Matches (Simera) | 100% Matches (Simera 2) |
| --- | --- | --- | --- | --- | --- |
| Close | 12 | 4744 | 18.0% | 20.0% | 20.0% |
| Close | 24 | 7309 | 6.6% | 38.1% | 42.9% |
| Close | 48 | 15822 | 3.5% | 18.4% | 15.8% |
| Distant | 12 | 2442 | 9.3% | 44.4% | 38.9% |
| Distant | 24 | 5093 | 4.6% | 35.6% | 26.4% |
| Distant | 48 | 11018 | 2.4% | 32.7% | 28.8% |

## 6 Discussion

The model presented in Section 4 provides an accurate representation of PCR. It has few parameters and assumptions and the output is shown to reflect real experimental results. The drawbacks of this model are related to speed limitations associated with the implementation of the algorithm and its usage is restricted to very small datasets which mean that it can’t be used for the majority of analyses.

A second model is presented in Section 5, the algorithm of which solves the problems associated with Model 1. Furthermore, the results obtained from simulations utilising the second model’s methodology show no indication of being any less accurate than those obtained from simulations involving the first model’s methodology.

The abundance and nucleotide composition of chimeras generated, both in *vitro* and *in silico,* can vary somewhat even when PCR is performed under the same conditions on identical samples. This is, presumably, caused by the random occurrences of PCR extension failure and makes it difficult to determine the degree of realism present in the data output from the simulations. However, results show that around a quarter of the chimeras found in the experimental data involving pooled nematode samples were reproduced perfectly during the simulated experiments and around a half were reproduced at a level of better than 99% similarity. Because of the potential for PCR and sequencing noise, it is reasonable to suggest that some of the closely matching chimeras were, in fact, exact matches.

The fact that many chimeras were reproduced exactly is encouraging, as are the results showing that the chimera break points are distributed similarly in experimental and simulated datasets. This is evidence that the chimeras are being generated in the same way, i.e. that the same ‘type’ of chimeras are being produced, even if their nucleotide composition is subject to natural variance. Furthermore, the distributions of break points on chimeras generated using the Simera algorithms compare favourably to the distributions of those on chimeras generated using Grinder. This is evidence that Simera generates more realistic chimeras than the best existing PCR simulators.

Section 1.1 discusses some existing PCR simulation software and notes that most available tools involve the selection of amplicons from a reference database by matching primer sequences to areas of similarity on the reference sequences. Because both Simera algorithms require ready-made amplicons as input, the models presented in this article work best when used in conjunction with existing software. For example, amplicons can be selected from a reference database using Primer Prospector and these amplicons can then be used as input for one of the Simera algorithms.

Overall, it can be concluded that both models presented in this article can be used to produce realistic simulated PCR output, particularly with respect to the chimeras generated during the process. In addition, the Simera 2 algorithm can be implemented sufficiently well to allow these simulations to be carried out on large, realistic datasets.

